# Neural codes in early sensory areas maximize fitness

**DOI:** 10.1101/2021.05.10.443388

**Authors:** Jonathan Schaffner, Philippe N. Tobler, Todd A. Hare, Rafael Polania

## Abstract

It has generally been presumed that sensory information encoded by a nervous system should be as accurate as its biological limitations allow. However, perhaps counter intuitively, accurate representations of sensory signals do not necessarily maximize the organism’s chances of survival. To test this hypothesis, we developed a unified normative framework for fitness-maximizing encoding by combining theoretical insights from neuroscience, computer science, and economics. Initially, we applied predictions of this model to neural responses from large monopolar cells (LMCs) in the blowfly retina. We found that neural codes that maximize reward expectation—and not accurate sensory representations—account for retinal LMC activity. We also conducted experiments in humans and find that early sensory areas flexibly adopt neural codes that promote fitness maximization in a retinotopically-specific manner, which impacted decision behavior. Thus, our results provide evidence that fitness-maximizing rules imposed by the environment are applied at the earliest stages of sensory processing.

## INTRODUCTION

One of the main goals of neuroscience is to understand what general principles explain the solutions evolution has selected to extract and process information from the environment. Half a century ago, it was postulated that sensory neurons should represent the sensory world as accurately and efficiently as possible by exploiting information about the statistical regularities of the environment, an idea known as *efficient coding* (1, 2). This principled approach has played a fundamental role in neural process theories in early sensory systems (3–7), and also in higher cognitive functions (8–14).

Efficient coding in sensory perception is typically assumed to be based on an information maximization criterion, that is, the sensory world must be represented as accurately as possible. One may think that this criterion makes sense for early sensory systems as this is precisely the role of a sensor: a good measurement instrument must reliably measure the environmental variable that it was built for. However, the information maximization criterion does not necessarily take into consideration the behavioral needs of the organism (15–19).

In line with this idea, there is evidence showing that early sensory systems represent not only information about physical sensory inputs, but also non-sensory information according to the requirements of a specific task and the behavioral relevance of the stimuli (20–24). However, this line of research provides no indication of the actual benefit of having such mixed neural representations at the earliest stages of sensory processing, or how this information could be used to efficiently guide behavior. Here, we provide a formal justification for these intriguing observations by testing the following hypothesis. Given that noisy communication channels always lose information during transmission, the brain will adapt to the fitness-maximizing rules of a particular environment at the earliest stages of sensory processing. Based on normative theoretical principles, we demonstrate that early visual structures in both insects and humans do indeed follow fitness-maximizing coding schemes.

## RESULTS

### Neurons in the blowfly retina code for contrast using a fitness maximization scheme

In a first attempt to test the hypothesis that the earliest stages of sensory processing incorporate fitness-maximizing encoding rules, we studied neural responses of retinal neurons in the blowfly—the large monopolar cells (LMCs)—which encode sensory information about visual contrast levels. To date, such neural codes are considered the first demonstration of efficient coding in biological organisms (25).

Visual features such as shape, color and patterns are important sensory signals that insects use to discriminate between competing flowers and fruit species, with visual contrast seeming to play a key role (27, 28). We propose that blowflies use knowledge of the different levels of contrast displayed by flowers and fruits in order to select food sources that promise more beneficial nutrients (i.e., reward). In other words, there is an association between contrast and reward that makes some contrast discrimination mistakes more costly than others. If this is the case, then a neural code that simply maximizes information accuracy in the LMCs would not maximize the fitness of the organism.

Concretely, we studied the following problem. Suppose that the the distribution of contrasts encountered by the blowfly in its natural environment is given by *f* (*s*). We define the function that transforms the contrast stimulus input *s* to neural responses *r* in the blowfly retina as *r* = *h*(*s*). Then, what is the optimal neural response shape *h*(*s*) if, given biological limitations, such a function can only generate a limited set of neural responses? Under this formulation, the following problem can be studied (29): find the optimal neural response function under two evolutionary optimization criteria, the *probability of mistakes* minimization criterion, and (ii) the *expected reward loss* minimization criterion.

To solve this problem, we assume that the organism must make choices between alternatives drawn from the stimulus distribution, *f* (*s*), which describes the relative availability of the different alternatives in its environment (e.g. how often a blowfly encounters a particular flower). The goal is to select the alternative that promises more reward to the organism, as this should lead the organism to maximize its fitness (15, 29). In the case of the blowfly LMCs, one may suppose that the blowfly must often make fine discriminations and that different contrast levels are linearly (and locally) related to different reward values.

If the goal of the organism is to minimize the number of erroneous responses (i.e., maximize discrimination *accuracy*), the optimal neural response *h*(*s*) matches the CDF of the stimulus distribution (Fig. 1b; see equation 5, Methods). However, this accuracy maximization strategy does not provide a precise account of the distribution of neural responses in the blowfly (Fig. 1b,c).

**Fig. 1.**
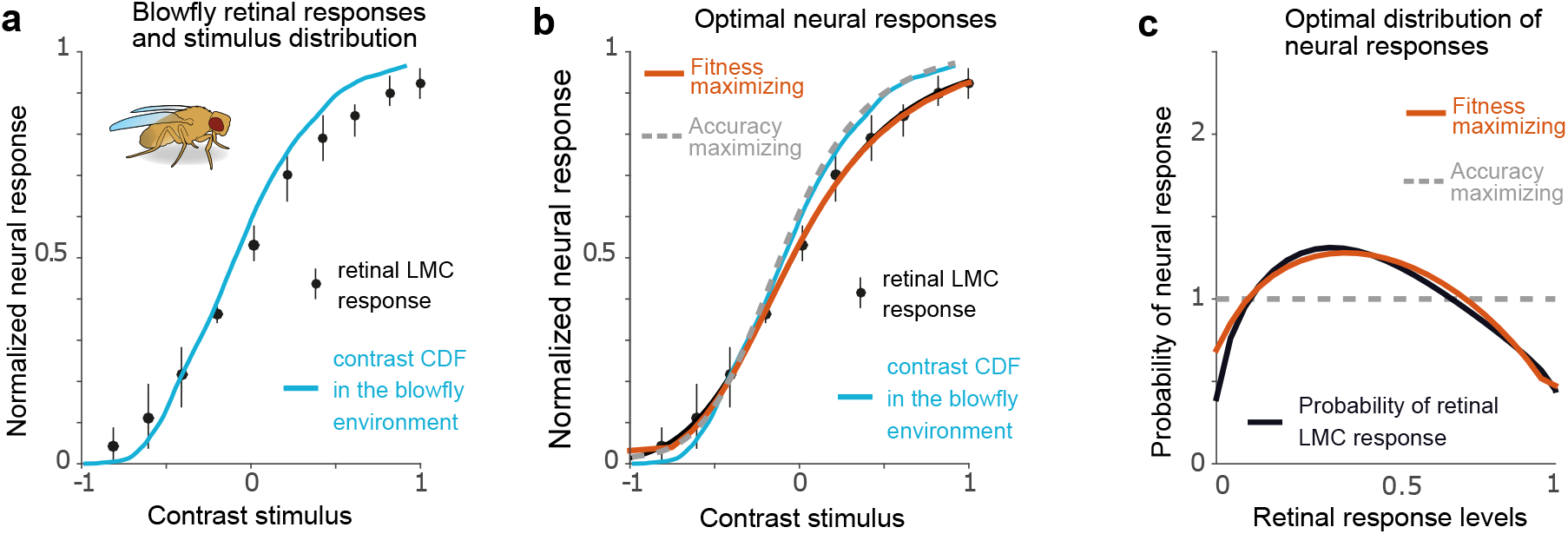
Blowfly LMC responses are better explained by fitness than information maximization coding schemes. **a)** Responses measured from the LMCs (black dots) and the cumulative distribution (CDF; blue line) of contrasts in the natural environment of the blowfly (25). If accurate perception of the environment is maximized by the LMCs, then the line indicating the CDF should lie directly on top of the dots indicating the empirical data. **b)** The black dots represent the same empirical data and the blue line the same contrast stimulus CDF as in panel a. The grey dashed line represents the predicted response function from an accuracy maximization code, but the orange line indicates a coding rule that maximizes fitness and matches the data better than the grey line. **c)** Neural response probability density distributions predicted by a fitness maximization rule (orange) also align better with the empirical data (black) than those predicted by infomax coding (dashed grey). This suggests that the fitness maximization model describes the empirical data more accurately than the accuracy maximization model. Interestingly, the same fitness-maximizing solution emerges when studying the *L*_*p*_ reconstruction error penalty, with optimal solution *p* = 0.5, which is the error penalty that best describes the LCM neural response data (26).

On the other hand, if the goal of the organism is to maximize the amount of *reward* received after many decisions, the optimal fitness-maximizing neural response *h*(*s*) provides a nearly perfect account of the neural responses of the blowfly retina (Fig. 1b,c; solution given in equation 6 in Methods). We emphasize that the remarkable overlap between the fitness-maximizing predictions and the empirical neural responses presented in Figure 1 are not the product of curve fitting, but instead these predictions emerge from the normative decision model, which has no degrees of freedom (Methods).

A common approach adopted in computer science and neuroscience to study the way in which a system penalizes estimation mistakes to optimize performance is via the *L*_*p*_ loss function defined as | *ŝ* (*r*) − *s* | ^*p*^, where *ŝ* is the sensory estimation, *s* is the true signal, and *p* determines how errors are penalized. A recent parameter estimation study asked what type of error penalty best explained the LMC data (26). However, in this study, the authors did not have explicit hypotheses for the potential evolutionary and behavioral meaning of different values for error penalties, and instead relied on numerical estimations of *p* that best explained the data. Remarkably, we demonstrate that the error penalty that provides a nearly perfect fit to the LMC response function corresponds to the blowfly LMC encoding function that guarantees maximal reward expectation to the organism (Fig. 1c, Methods).

### Adaptive fitness-maximizing sensory codes in humans

While the predictions of reward-maximizing sensory codes and the blowfly LMC data show a striking similarity (Fig. 1), it remains only anecdotal evidence for our hypothesis that early sensory areas *adapt* to the organism’s behavioral needs. We don’t know the specific function linking contrast to fitness for the blowfly, and we can’t show that the code used in their retinas adapts between contexts. Therefore, we implemented an experiment that allows us to test whether fitness-maximizing codes are not only present, but also flexibly adaptive in early visual areas in humans.

To date, it has been widely accepted that the default neural code for orientation perception in humans is information maximization (infomax) coding (30). This code will minimize the probability of mistakes. One reason that infomax coding may typically explain human perception in regard to orientation well is that, for humans, orientation information does not typically signify reward and is instead used for navigation purposes. Thus, the fitness maximizing code for orientation perception may be equivalent to infomax under standard conditions. Our experiments create a context that deviates from these standard conditions to test if human early visual areas adapt in a manner predicted by the theory of fitness maximization. In the experiments with human participants, we take advantage of the fact that neurons in the retina and early visual areas have receptive fields that include specific subsets of the visual field. This feature of the visual system, in conjunction with computational modeling techniques, allows us to test our hypotheses without needing to measure neurons directly.

We designed a behavioral task in which, on any given trial, human participants had to choose which of two simultaneously presented orientation stimuli, *s*_1_ or *s*_2_, was more diagonal (i.e. closer to a 45-degree angle; Fig. 2, Fig. 3a). The key manipulation in our task is that decisions were made in two different stimulus-reward association contexts. In one context, participants were paid a fixed reward for correct discrimination of the more diagonal stimulus in each trial, and received no reward for incorrect decisions (henceforth accuracy context *K*_acc_). In the second context, participants were rewarded depending on the stimulus *s* they selected in each trial, and the amount of reward was linearly mapped to the degree of “diagonality” of the input stimulus (henceforth reward context *K*_rew_). Crucially, the prior distribution of sensory signals *f* (*s*) was exactly the same in both contexts. Stimuli close to cardinal orientations were presented most often to match the statistics of natural scenes that humans typically encounter (31) (Fig. 3a). This experimental design allows us to test the competing hypotheses that neural codes in early sensory areas (i) maximize accurate representations of the environment and are thus constant in both reward contexts, or (ii) adapt between contexts in order to instantiate efficient coding strategies that maximize fitness.

**Fig. 2.**
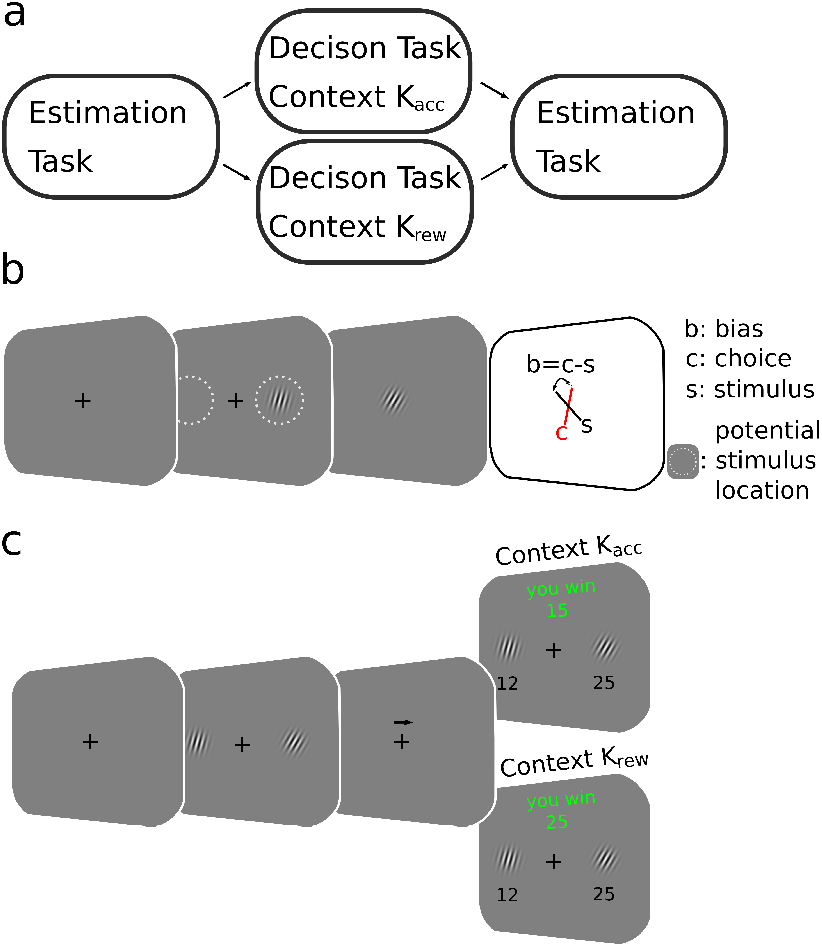
Behavioral paradigm in human participants. **a)** Participants performed the “Estimation task” before and after training in the “Decision task”. They completed the Decision task either in the reward context, *K*_rew_, or the accuracy context, *K*_acc_ (see c below). **b)** Estimation task. After perceiving a Gabor patch stimulus on the left or right side of the screen, participants had to rotate a Gabor cue in the middle of the screen until its orientation matched the orientation of the perceived stimulus. **c)** In the Decision task participants decided which of two Gabor patch stimuli is more diagonal. In context *K*_acc_, participants received a constant reward for a correct decision, whereas in context *K*_rew_, reward magnitude was linearly related to the degree of diagonality of the stimuli.

**Fig. 3.**
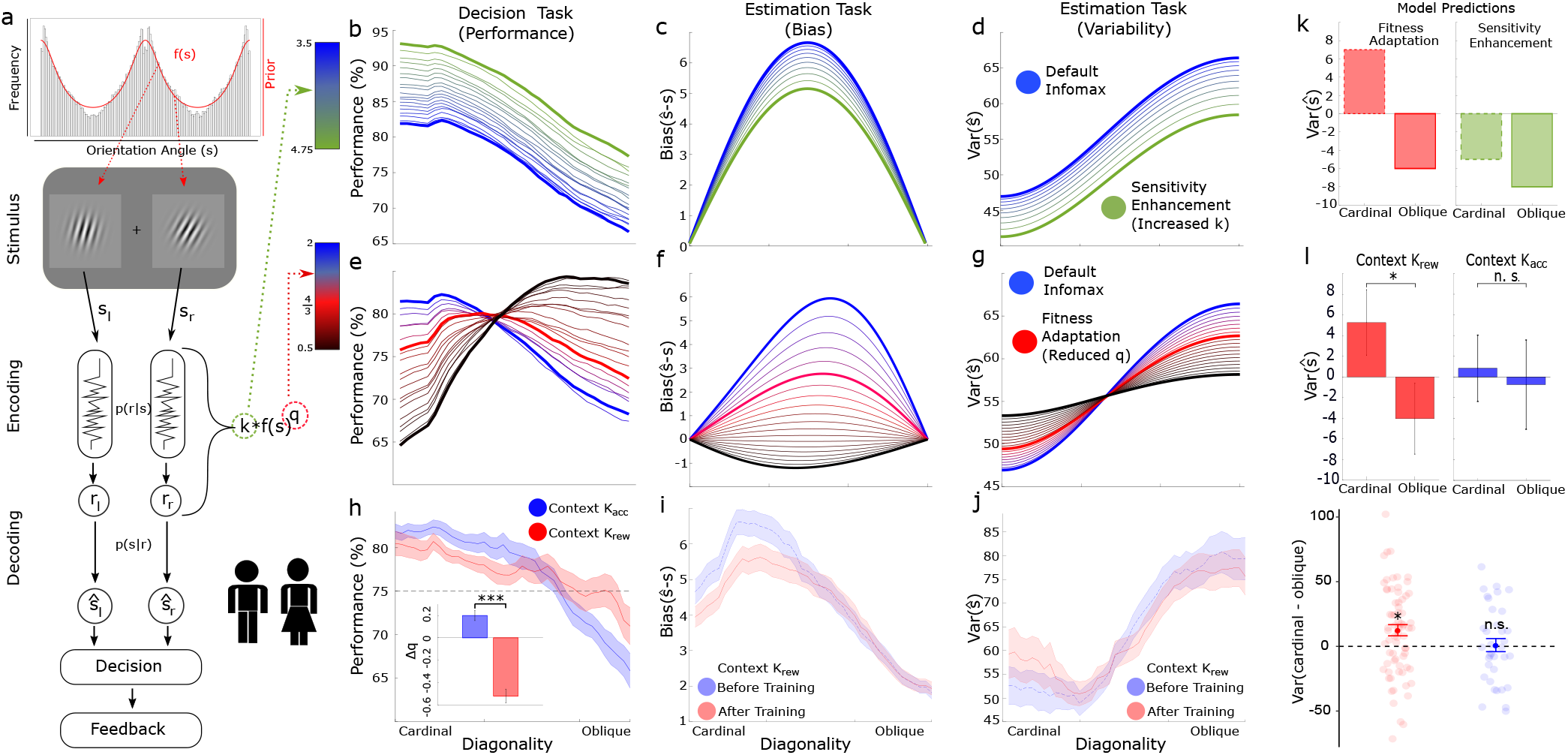
Inference model, theoretical predictions, and empirical results. **a)** Simplified schema of the decision task. The orientation of the Gabor patches was drawn from the distribution of edges in natural scenes. The orientation *s* of the perceived Gabor patches is encoded with the internal response *r*. The corresponding likelihood function *p*(*r*|*s*) is constrained by the encoding rule. The prior *f* (*s*) is combined with the likelihood to generate an estimation *ŝ*. The encoding rule depends on the model parameters *q* and *k*. Participants made decisions in two different stimulus-reward association contexts. In context *K*_acc_, participants were paid a fixed reward for correct discrimination of the more diagonal stimulus in each trial. In *K*_rew_ context, participants were rewarded depending on the stimulus *s* they selected in each trial, and the amount of reward was linearly mapped to the degree of “diagonality” of the input stimulus (see Figure 2). Model predictions for the “Decision task” assuming enhanced sensitivity *k*. The blue line represents a common infomax encoding model. As sensory precision increases, performance increases consistently at all orientations (greener lines). **c**,**d)** Bias and variability predictions in the “Estimation task”. Increasing sensitivity *k* decreases estimation bias (*ŝ*−*s*) (panel c), and generally reduces the variance over the whole range of diagonality (panel d). **e)** Performance predictions as a function of power-law encoding *q*. As in panel b, the blue line represents the infomax model. The thick red line shows the prediction of the fitness-maximizing model. As *q* decreases, performance drops for the more cardinal trials, and increases for the more oblique trials. **f)** If *q* decreases for a constant capacity level, estimation biases should decrease. **g)** As *q* estimation variability decreases for oblique angles, it increases for cardinal angles. **h)** Empirical performance in the “Decision task”. The light blue line shows participants in the accuracy context, the red line in the reward context. The difference between contexts matches the predictions for the fitness adapting model. The inset shows the change in *q* after participants engage in the accuracy (blue) or reward (red) context. Again, *q* changes as expected by a fitness-maximizing adaptive code. **i)** The results of the “Estimation task” show a significant reduction of the bias for all sessions with training in context *K*_rew_. **j)** The change of the variance is consistent with a fitness-maximizing code as seen in panel g. For sessions in context *K*_rew_, variance decreases for oblique stimuli and increases for cardinal stimuli. **k)** Prediction of estimation variance changes for cardinal and oblique angles. Variance generally decreases for increased sensitivity (green bars). However, if fitness-maximizing adaptation takes place, then variance should increase for cardinal angles and decrease for oblique angles. **l)** In line with these predictions, in *K*_rew_ context (red), estimation variance increases for cardinal edges (*<* 11.25°) and decreases for oblique edges (*>* 33.75°). There is no significant change in the *K*_acc_ context (blue). The lower plot shows the difference in variance between oblique and cardinal orientations at the individual level (red = *K*_rew_, blue = *K*_acc_). Each dot represents a participant and the darker dots show the group mean. Error bars denote SE in all panels. (* *P*_MCMC_ *<* 0.05, *** *P*_MCMC_ *<* 0.001).

We employed a general method for defining efficient codes by investigating optimal allocation of limited neural resources (32). Based on this framework, sensory precision should be proportional to the amount of resources available *k* and the prior distribution *f* (*s*) raised to a power *q*, hence known as the power-law efficient code, *k ·f* (*s*) ^*q*^. We show that an advantage of employing this framework is that there is a direct link between the power-law efficient codes and the fitness maximization solutions for the contexts that we consider here (Methods). In brief, the connection of the power-law efficient codes with accuracy vs reward maximization objectives is the following: If the power-law parameter *q* is relatively low, it shades some density away from where *f* (*s*) is high and relocates it where it is low. The reason for this spreading of coding resources is that, even though observing *f* (*s*) is unlikely, and thus the probability of error will be low, if there is an error, it could be a very costly mistake. Therefore, when stimuli are directly associated to rewards relative to situations in which all mistakes are equally costly, it pays to allocate more neural resources to the segments of the stimulus space where *f* (*s*) is low (Fig. 3e and Figure S1). This theoretical link allowed us to derive various qualitative predictions that we used to test whether humans indeed adopt fitness-maximizing, as opposed to information-maximizing, neural codes in early sensory areas.

A first prediction is related to sensory discrimination differences in the two reward association contexts considered here. If participants maximize fitness under limited resources, discrimination accuracy should be higher for cardinal orientations in *K*_acc_ than in *K*_rew_, whereas for diagonal (i.e., oblique) orientations discriminability must be higher in *K*_rew_ than in *K*_acc_ (Fig. 3e). In line with the theory, we found that participants performed better in trials showing cardinal orientations in the context *K*_acc_ relative to *K*_rew_, and this pattern reversed for trials showing more diagonal orientations (interaction *s* \**K*: *β* = 0.26 ± 0.11, *P*_MCMC_ = 0.011). Despite these relative differences, the fitness maximization theory predicts that, for both optimization objectives, discriminability should be higher for regions of the stimulus distribution prior with higher density (i.e., cardinal orientations) given that these stimuli occur more frequently in all contexts. Our data are consistent with this prediction as well (main effect of orientation *s*: *β* = 0.48 0.07, *P*_MCMC_ *<* 0.001). Note that these results are not a product of increased sensitivity due to task experience, given that increased sensitivity would result in a general improvement in task performance across the whole orientation space (Fig. 3b), which does not account for the significant interaction effects that are present in our data (Fig. 3h) and predicted by the fitness-maximizing model (Fig. 3e). These results provide support for our hypothesis that perceptual coding of sensory information adopts a fitness-maximizing code.

### Signatures of fitness-maximizing adaptation at sensory estimation stages

We sought to determine whether the form of efficient adaptation observed in the decision task only takes place in downstream decision circuits, or if it is already implemented at earlier processing stages, for instance, in the circuits that generate estimations of sensory stimuli.

To answer this question, we investigated a second prediction of the theory, which can be derived from orientation estimation biases and their relation to the oblique-effect illusion during orientation estimation tasks (30). We asked participants to perform an edge orientation estimation task before and after the contextual decision-making task (Figure 2, Methods). This allowed us to determine whether efficient adaptation occurs at the level of sensory estimations, and whether the neural codes follow a fitness-maximizing or infomax scheme after participants were exposed to context *K*_rew_. Departing from the assumption that for orientation-tuned neurons the default strategy is an infomax code, fitness maximization predicts that, after participants adapt to context *K*_rew_ in the decision task, the magnitude of repulsive estimation biases away from the prior distribution should decrease (Fig. 3f). In line with the fitness maximization predictions, we found that repulsive biases were smaller after exposure to context *K*_rew_ (Fig. 3i; interaction time(after-before): *β* = −0.37 ± 0.18, *P*_MCMC_ = 0.023. In contrast, the fitness maximization applied to context *K*_acc_ predicts that there should be no change in estimation biases. In line with this prediction, we found no significant change in estimation biases after *K*_acc_ training (interaction time(after-before): *β* = +0.42 ± 0.32, *P*_MCMC_ = 0.91). Moreover, consistent with fitness maximization, but contrary to strict infomax coding, we found that the decreases in repulsive biases were greater following *K*_rew_ than *K*_acc_ (interaction *K\**time(after-before): *β* = −0.79 ± 0.32, *P*_MCMC_ = 0.0085).

A third prediction is related to the trial-to-trial fluctuations (variability) in the estimation of sensory stimuli. The fitness maximization hypothesis predicts that after participants adapt to context *K*_rew_ in the decision task, estimation variability for input stimuli near cardinal orientations will be slightly higher. However, there should be lower variability for more diagonal orientations (Fig. 3g,k). This is because the theory predicts a shift in coding resources from high probability cardinal orientations to low probability diagonal orientations. Once again, our data are in line with these theoretical predictions. We found that after *K*_rew_ training, variability in sensory estimations was higher for cardinal orientations and lower for diagonal orientations (*β* = −2.7 ± 1.49, *P*_MCMC_ = 0.035; Fig. 3j,l). Crucially, fitness maximization predicts that after exposure to context *K*_acc_, there will be no change in estimation variability for cardinal or diagonal orientations. In line with this prediction, we found no significant change i n the variance after *K*_acc_ training (*β* = −0.8 ± 2.17, *P*_MCMC_ = 0.36, Fig. 3l). We also find an interaction trend between context (*K*_rew_ vs *K*_acc_) and cardinality (interaction *K* *cardinality: *β* = −8.75 ± 5.4, *P*_MCMC_ = 0.07); the robustness of this result was confirmed by the analysis of individual differences presented in the next paragraph and, subsequently, in the independent sample in Experiment 2.

### Signatures of fitness-maximizing a daptation a re related to efficient neural recoding and not to sensitivity enhancement

Next, we investigated the alternative hypothesis that the bias and variance changes observed after adaptation to context *K*_rew_ were caused by a general enhancement in sensitivity *k* due to practice effects. This hypothesis predicts a reduction in estimation bias (Fig. 3c) that is similar to the qualitative prediction from the fitness-maximizing hypothesis (Fig. 3f). However, the predictions of the two hypotheses differ with regard to estimation variability. Enhanced sensitivity predicts a general decrease in variability across the entire range of cardinality space (Fig. 3d), whereas the fitness-maximizing hypothesis predicts the trade-off between cardinal and diagonal orientations described above (Fig. 3g). Accordingly, our data (Fig. 3j) are consistent with the fitness-maximizing hypothesis (Fig. 3g), and not the enhanced sensitivity hypothesis (Fig. 3d). In order to further substantiate this conclusion, we tested if the amount of change in the sensory estimations that participants showed after contextual *K*_rew_ decisions was associated with the resource reallocation of neural resources (parameter *q*). To this end, we fit the encoding model to the choice data to estimate parameters *q* and *k*. In line with the theory, we found that in the context *K*_rew_ the value *q* decreased between the first and last part of the decision experiment by Δ*q* = −0.5225 ± 0.062, *P*_*MCMC*_ *<* 0.001 (Fig. 3h). Comparing the change in *q* (i.e., Δ*q*) between the two contexts showed that there was a significantly greater decrease in *K*_rew_ than *K*_acc_ (Δ*q* = −0.7206 ± 0.08, *P*_*MCMC*_ *<* 0.001, Fig. 3h). Additionally, in line with the fitness-maximizing hypothesis, we found that changes in the parameter *q* during the decision task were significantly related to estimation bias changes in the sensory estimation task (*β* = −24.67 ± 11.01, *P*_MCMC_ = 0.015), controlling for changes in sensory precision *k*. Circularity is not an issue in this analysis because the encoding model parameters *q* and *k* result from fits to the independent data from the choice task, and the estimation task data were acquired before and after the decision task (Fig. 2). Taken together, our results clearly indicate that the empirically observed behavioral changes in the estimation of sensory signals were not caused by simple practice-related sensitivity enhancement, and further support the fitness-maximizing coding hypothesis.

### Fitness-maximizing codes are retinotopically specific

A key conclusion that we draw from our results is that fitness-maximizing adaptation in both the decision task (Fig. 3h) and the estimation task (Fig. 3i,j) appears to have a common origin that does not depend on comparisons between decoded stimuli in downstream decision circuits. To explicitly test if fitness-maximizing neural codes are indeed present at the earliest stages of sensory processing, we conducted a variation of our first experiment in an independent sample. In this modified estimation task, participants were presented with an orientation stimulus in one of four spatial locations. Crucially, participants were trained in only two of these locations in the *K*_rew_ decision task (Fig. 4a). If adaptation is retinotopically specific, then changes in bias and variance (after vs before contextual training) should be specific to the retinotopic locations trained during the decision task. In line with the fitness-maximizing predictions, we found that reductions in estimation bias were present in the trained locations (*β* = −0.60 ± 0.24, *P*_MCMC_ = 0.0078). There was no significant bias reduction in the untrained locations (*β* = −0.05 ±0.24, *P*_MCMC_ = 0.43). Furthermore, the effect was significantly greater in the trained than un-trained location (interaction location*time(after-before): *β* = −0.55 ± 0.32, *P*_MCMC_ = 0.0425; Fig. 4b). We also found the pattern predicted by a fitness-maximizing code in the location specific changes in estimation variability (interaction location*time(after-before): *β* = −15.8 ± 6.4, *P*_MCMC_ = 0.0078; Fig. 4c). Together, these results confirm the retinotopic specificity of fitness-maximizing coding rules.

**Fig. 4.**
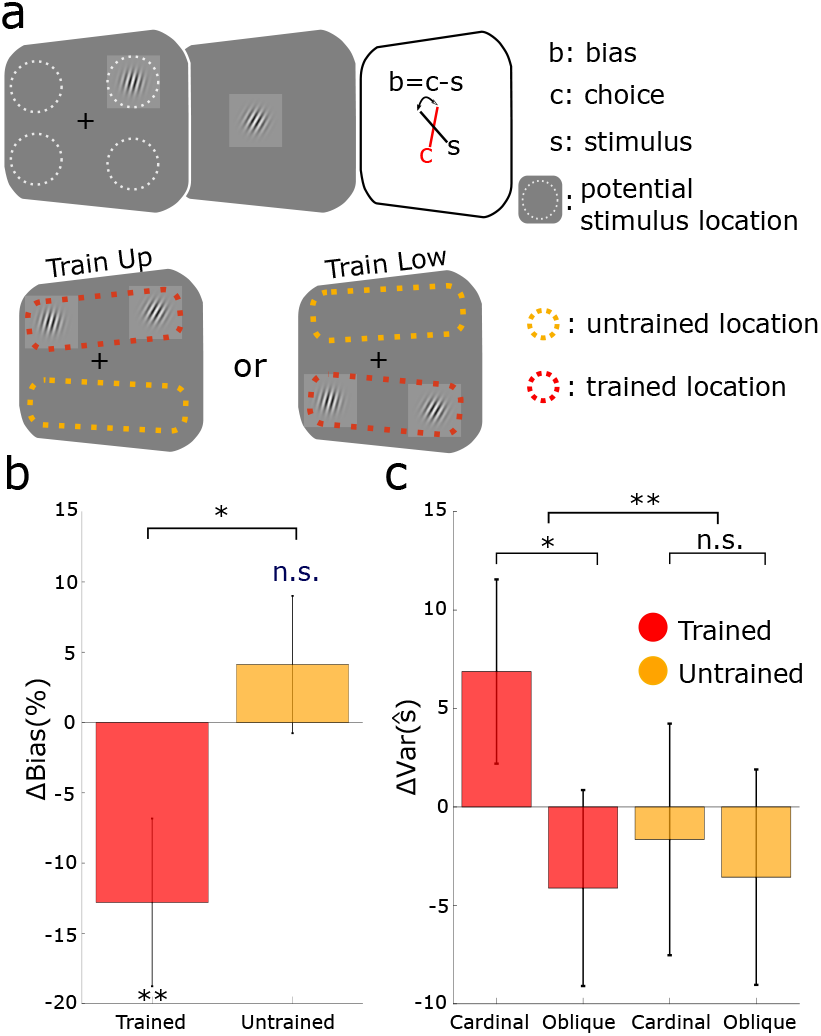
Retinotopic Specificity experiment. **a)** In a variation of Experiment 1, Experiment 2 contained four possible locations for stimulus presentation. However, only two of these locations were trained in the “Decision task”. In each session observers were only trained in the upper or lower locations. In the plots, the red shading indicates the trained locations, and orange shading the untrained locations. **b)** Proportion change in estimation bias for the trained and the untrained locations. The bias only decreased in the trained location. **c)** Change in estimation variability for the trained and the untrained locations. For the trained locations, variance changes followed the fitness-maximizing predictions for cardinal (*<* 11.25*°*)) vs oblique (*>* 33.75*°*) stimuli. This variance change was specific to the trained location. Each dot represents an individual participant and the darker dots indicate the group mean. Error bars denote SE. (* *P*_MCMC_ *<* 0.05, ** *P*_MCMC_ *<* 0.01).

## DISCUSSION

Our theoretical and empirical results provide evidence that the earliest stages of sensory processing have evolved to encode environmental stimuli in order to promote fitness maximization, and not necessarily to maximize perceptual accuracy. We have shown this to be the case in both insects and humans, which indicates that fitness-maximizing perceptual coding schemes are present in early sensory systems across taxa. Crucially, our findings indicate that down-stream circuits do not necessarily need to continuously compute reward distributions based on stimulus-outcome associations because this information can be efficiently embedded in the neural codes of early sensory systems. This notion is supported by a recent study showing that functional remapping of stimulus-reward contingencies in early sensory areas causally depends on top-down control signals from prefrontal structures (23, 24, 33). We argue that this gives the organism the advantage of rapidly transmitting behaviorally-relevant information encoded by early sensory systems to downstream circuits specialized in action, learning, or decision-making.

Beyond the obvious relevance for biological organisms, our results may have important implications in ongoing developments in artificial intelligence (AI) as well. Recent deep generative models show a remarkable ability to encode high-dimensional signals into latent factors under the objective of accurately predicting the local environment with specific encoding constraints. However, based on our results, such an optimization objective will not necessarily match those present in biological organisms. Interestingly, a recent successful AI model (34) proposed instead that representation formation should be driven by the need to accurately predict the motivational value of experiences. Our results validate this notion, and imply that the development of AI algorithms that aim to resemble neurobehavioral function should go beyond the objective of maximizing only the accurate transmission of information and account for the motivational aspects of the environment that enable the organism (or the artificial agent) to maximize fitness.

Finally, although drawn from a different domain of behavior, our results lend substantial support to economic theories positing that context-dependent utility functions should maximize expected reward rather than the expected accuracy of decisions guided by reward (15, 29, 35). The corroborating evidence presented in our work grounded on the principles of neural coding and decision behaviour should help to advance the development and refinement of these theories within economics and related disciplines of evolutionary biology and social sciences (36–38).

## Methods

### Participants

Participants were recruited by the Center for Neuroeconomics at the University of Zurich, Switzerland (n=81 in total, n=36 in Experiment 1, n=45 in Experiment 2). Participants were instructed about all aspects of the experiment and gave written informed consent. None of the participants suffered from any neurological or psychological disorder or took medication that interfered with participation in our study. Participants received fixed monetary compensation for their participation in the experiment, in addition to receiving variable monetary reward depending of task performance (see below). The experiments conformed to the Declaration of Helsinki and the experimental protocol was approved by the Ethics Committee of the Canton of Zurich.

Participants who failed to follow the eye fixation instructions on more than 25% of trials were excluded from the data analysis (n=12). We measured the performance of participants in the training tasks and excluded participants that were unable to perform the task at the easiest difficulty level (n=11). Additionally, we had to exclude three participants due to technical problems with the data collection. Thus, the final sample comprised n=55 participants (n=25 in Experiment 1, n=30 in Experiment 2).

### Experimental design and stimuli

Stimuli were generated with Matlab, using the Psychtoolbox and displayed on a screen using the following experimental protocol. Participants were sat one meter from the screen. The angle of the head was kept stable with a chin rest. The height of the chin rest was adjusted to position the center of the screen at the height of the eyes. As orientation stimuli, we used oriented Gabor patches, presented on a grey background. Each patch was composed of a high-contrast 3 cycles/degree sinusoidal grating convoluted with a circular Gaussian with width 0.41° and subtended 2.98° vertically and 2.98° horizontally. In experiment 1, all Gabor patches were presented so that the centers fell 5.7° to the left or right of the center of the monitor and on the horizontal midline. In experiment 2, the Gabor centers fell 4.7° to the left and right of the vertical midline, and 4.7° above or below the horizontal midline.

#### Eye Tracking

Eye-tracking data was acquired using an ST Research Eyelink 1000 eye-tracking system. Gaze position was sampled at 500 Hz. Eye movements away from fixation were computed for the window corresponding to stimulus presentation. For every saved position the absolute distance to the fixation cross was computed. If the absolute distance exceeded 4° of visual angle the trial was marked to include an eye movement. For most participants, the average number of trials with eye movements was less than 5%. Participants (n = 12) who made eye movements that exceeded 4° of visual angle on more than 25% of trials were excluded from all analyses.

#### Experiment 1

Participants performed the experiment in multiple sessions to allow for training within both contexts on different days. The order of the accuracy (*K*_*acc*_) and reward (*K*_*rew*_) context training was counter-balanced across participants. In total, every participant completed 240 trials in the “Estimation task” and 400 trials in the “Decision task”.

#### Experiment 2

Training in the “Decision Task” was performed either in the two upper locations or in the two lower locations. Participants were randomly allocated to one of the two conditions. For the estimation task trial locations were evenly distributed between all four possible locations. In total every participant completed 400 trials in the “Estimation task” and 360 trials in the “Decision task”.

#### Orientation estimation task

Before the start of every trial participants had to fixate on a cross in the middle of the screen. At the beginning of the trial an arrow appeared for 0.5 seconds to indicate on which side the stimulus would be shown. Afterwards, the stimulus appeared on the indicated side for 0.6 seconds. The orientation of the stimulus was determined randomly from (0 − 179°). During stimulus presentation, the participant had to continue fixating on the cross. After the stimulus disappeared, a Gabor patch appeared in the middle of the screen. The participant could, by pressing and holding the left mouse button, rotate the new Gabor patch until its perceived orientation matched the orientation of the previously observed target stimulus. The participant could end the trial by pressing the space key. After five seconds the trial ended automatically. Trials were separated by a random 1.5-2 seconds intertrial interval.

#### Binary judgment (Decision-making) task

The fixation cross turned black to indicate the start of a trial. After 0.5 seconds, two Gabor patch stimuli appeared. The orientation of one of the stimuli was drawn from the approximate distribution of edges in the real world (31). The orientation of the second stimulus was adjusted by a participant-specific difficulty score to keep performance at approximately 75% accuracy for all participants. On average the stimulus orientation followed a prior distribution *f* (*s*) described by Equation 1 and shown in Fig.3a.

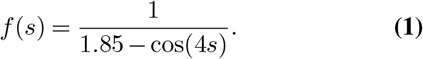

The stimuli were displayed for 0.6 seconds. During stimulus presentation participants had to fixate on the cross in the middle of the screen. When the stimuli disappeared, participants had 2.5 seconds to decide which stimulus was more oblique. Independent of the reaction time the full 2.5 seconds had to be waited out. Afterwards the 2 stimuli were shown again in their positions and the result of the choice, as well as the orientations of the stimuli, were displayed for 3 seconds until the trial ended. Trials were separated by a 1.5-2 second intertrial interval.

### Retinal large monopolar cells (LMCs) experiment in the blowfly

Here we provide a brief description of the data collected in Laughlin’s seminal work (25), which we reanalyze in this work. In order to derive the prior for the sensory stimulus of interest *f* (*s*), the researcher measured the distribution of contrasts that occur in woodland settings of the blowfly environment. This data was used to construct a histogram, which was later transformed to a CDF (Fig. 1a, Figure S1). Here we used this CDF to reconstruct the PDF *f* (*s*) (Figure S1). Once the prior distribution was obtained, the fly was placed in front of a screen with a light emitting diode (LED). At the beginning of each trial the LED luminance was set to the screen luminance and then changed to a new luminance drawn from the prior distribution *f* (*s*) for 100 ms. The stimulus *s* was defined as the proportional change of the difference between the background and LED luminances.

### Fitness-maximizing neural codes

In this section we provide a detailed description of the connection between (i) the *L*_*p*_ reconstruction error, (ii) the efficient code that maximizes reward expectation, and (iii) the power-law efficient codes briefly described in the main text.

Suppose that the stimulus distribution is given by *s* ∼ *f* (*s*). The function that transforms the input *s* to neural responses *r* is given by *r* = *h*(*s*). While the mapping *h*(*s*) is deterministic, here we assume that errors in neural response *r* follow a distribution *P* (*r* | *h*(*s*)). We apply a general approach that takes into consideration optimality criteria accounting for how well stimulus *s* can be reconstructed (*ŝ*) from the neural representations *r*. Wang and colleagues introduced a general formulation of the efficient coding problem in terms of minimizing the error in such reconstructions *ŝ* (*r*) according to the *L*_*p*_ norm as a function of the norm parameter *p* (39). In brief, the reconstruction is assumed to be based on the maximum likelihood estimate (MLE) of the decoder in the low noise regime, where *P* (*r* | *h*(*s*)) is assumed to be Gaussian distributed.

The goal is to find the optimal mapping function *h*^*\**^(*s*) to achieve a minimal *L*_*p*_ reconstruction error for a any given prior stimulus distribution *f* (*s*). More formally the problem is defined as: find *h*^*\**^(*s*) such that

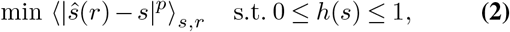

where, without loss of generality, we assume that the operation range of the neuron is bounded between 0 and 1. It is possible to show that the optimal mapping *h*^*\**^(*s*) is given by Eq. 3 (39).

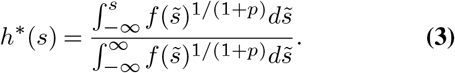

If we define

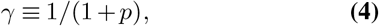

we observe that the normalized power function of the stimulus distribution *f* in Eq. 3 is the escort distribution with parameter *γ* (40). Note that under this framework, Infomax coding is given by the norm parameter *p* → 0, and therefore *α* = 1, thus leading to the result that *h*(*s*) is the CDF of the prior distribution.

#### Connecting efficient L_p_ error-minimizing codes and behavioral needs

We now turn to investigations from the field of Economics that studied the following problem: For a given distribution *f* (*s*) in the environment, what is the optimal shape of the subjective *utility function* (i.e., *h*(*s*)) if such function can only take a large but limited set of *n* discrete subjective values (i.e., the internal readings, *r*) that code for any given stimulus *s* (15, 29). The utility function is thus restricted to a set of step functions with *n* jumps, each corresponding to a utility increment of size 1*/n*. In this case, discrimination errors originate from the fact that the organism cannot distinguish two alternatives located at the same step of the utility function (note that this is different from the definition of the response *r* following a Gaussian distribution *P* (*r* | *h*(*s*)) in the neural coding problem defined earlier; however, it can be shown that these two problems lead to the same analytical solutions). Under this formulation, the following problem was studied: find the optimal utility function (*h*^*\**^) under two evolutionary optimization criteria, (i) the *probability of mistakes* minimization criterion, and (ii) the *expected reward loss* minimization criterion.

To solve this problem, we assume that the organism repeatedly makes choices between two alternatives drawn from the stimulus distribution *f* (*s*), where we may suppose that stimuli are linearly mapped to a reward value. The organism is endowed with a utility function that assigns a level of reward to each possible stimulus *s* from *f* (*s*). The alternative that promises more utility to the organism is chosen (29).

If the goal of the organism is to minimize the number of erroneous responses (i.e., maximize discrimination *accuracy*), the optimal utility function 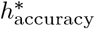 is given by

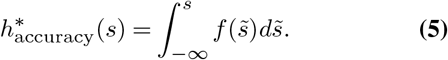

According to this solution, the power parameter of the escort distribution in Eq. 3 is given by *γ* = 1, which corresponds to the Infomax strategy.

On the other hand, if the goal of the organism is to minimize the expected reward loss (i.e., maximize the amount of *reward* received after many decisions) and stimuli are linearly mapped to reward value, the optimal utility function 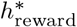 is given by

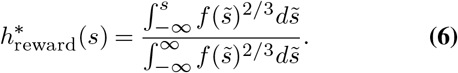

According to this solution, the power parameter of the escort distribution in Eq. 3 is given by *γ* = 2*/*3, which corresponds to optimizing the *L*_*p*_ minimization problem with parameter *p* given by

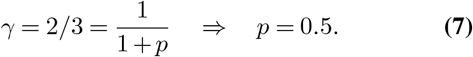

Remarkably, this normative fitness-maximizing solution is the error penalty that best describes the LMC data (26) (result reported in main text and Fig. 1). Additionally, please note that the solutions provided in equations 5 and 6 are derived based on maximizing the accurate choices and reward expectation, respectively, without any assumptions about maximizing information efficiency as a goal in itself.

#### Connection to power-law efficient codes

We employed a general method for defining efficient codes by investigating optimal allocation of Fisher information *J* given: (i) a bound of the organism’s capacity *c* to process information, (ii) the frequency of occurrence *f* (*s*), and (iii) the organism’s goal (e.g., maximize perceptual accuracy or expected reward). Here we employ a general method for defining efficient codes by investigating optimal allocation of Fisher information according to

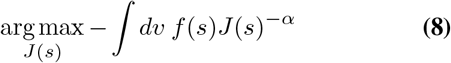

subject to a capacity bound

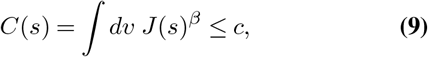

with parameters *α* defining the coding objective and *β >* 0 specifying the capacity constraint (32). The solution of this optimization problem reveals that Fisher information should be proportional to the prior distribution *f* (*s*) raised to a power *q*, which is therefore referred to as the power-law efficient code

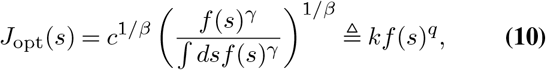

where *q* = 1*/*(*β* + *α*), and *γ* = *β/*(*β* + *α*). Note that power-law parameter *q* is multiply determined and to make progress in identifying it we need to make some further assumptions. Here we opted for setting *β* = 0.5, as previously proposed in the standard infomax framework (30), however our conclusions are not affected by the specific value of *β*. This means that *α* determines how Fisher information is allocated relative to the prior, influencing the values of both *q* and *γ*. Recall that the Infomax coding rule implies *γ* = 1 and therefore an efficient power-law code *q* = 2. Also recall that the reward expectation rule implies *γ* = 2*/*3 and therefore an efficient power-law code *q* = 4*/*3 (which in turn implies a *L*_*p*_ norm parameter *p* = 0.5). Thus, the power-law efficient codes allow us to establish a connection between behavioral needs in the contexts studied in this work (*K*_acc_ and *K*_rew_) and parameter *γ*, which incorporates the goals of the organism under the resource-constrained framework that we study here.

#### Optimal inference

When specifying an inference problem using such an encoding-decoding framework, a key aspect for generating predictions of decision behavior is to obtain expressions of the expected value and variance of the noisy estimations *ŝ* for a giving value input *s*_0_. However, we first need to specify the encoding and decoding rules. We adopt an encoding function *P* (*r* | *s*) associated with the power-law efficient code that is parameterized as Gaussian (32)

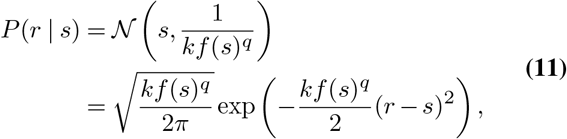

and therefore Fisher information is allocated using a *s*-dependent variance *σ*^2^ = 1*/kf* (*s*)^*q*^. While we are aware that in our study the stimulus space is circular, given that discriminability thresholds are relatively low for orientation discrimination tasks in humans, it is safe to assume that the likelihood function can be locally approximated as a Gaussian distribution.

At the decoding stage, the observer computes the posterior using Bayes rule

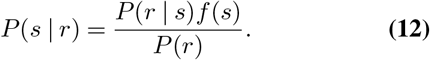

Theoretical and empirical evidence suggests that for orientation estimation tasks, estimates are typically biased away from the prior. This suggests that humans employ an expected value estimator of the posterior, at least for the infomax case (30).

The expected value of the estimator can be defined as the input stimulus *s*_0_ plus some average bias *b*(*s*_0_). Using analytical approximations under the high signal-to-noise regime, it is possible to show that the bias for the posterior expected value estimator can be approximated by

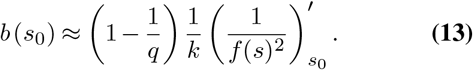

In a previous study, using model simulations and exploring parsimonious functional forms, it was shown that the proportionality constant of the bias term can be approximated by (32)

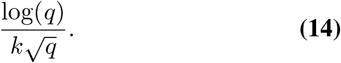

It can be shown that the analytical solution and the simulation-based solution of the proportionality constant are approximately equivalent for a range of *q* values relevant to our work (e.g., *q ∈* [0.5, 2]), that is

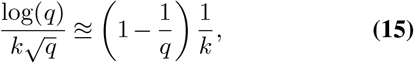

thus validating the results derived in the analytical approximations that we used in the current work. However using either function does not affect the qualitative or quantitative results in our study.

Using this result, the expected value of the estimators is given by

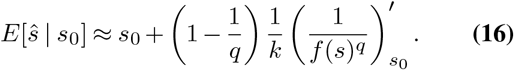

As already defined in the description of the behavioral task, in this study, we used a parametric form of the prior that closely resembles the shape of the natural distribution of orientations in the environment (31)

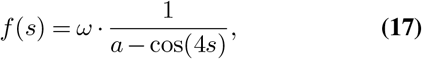

with *a >* 1 determining the elevation (steepness) of the prior, and *ω* a normalizing constant. Using this parameterization of the prior, we can obtain an explicit analytical approximation of the bias

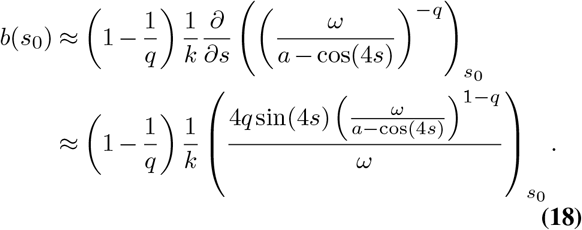

We can also obtain an analytical approximation of the variance under the high signal-to-noise regime using the Cramer-Rao bound formulation

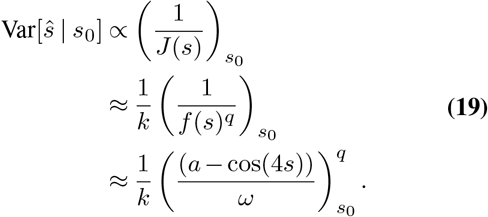

Thus, we can use equations 18-19 to derive the predictions presented in Figs. 2b-g.

Finally, assuming that the estimators are normally distributed using the expected value and variance derived above, the probability that an agent chooses an alternative with orientation value *s*_1_ over a second alternative with orientation value *s*_2_ (recall that in our experiment the decision rule (objective) of the participants is to choose the orientation perceived as closer to the diagonal orientation) is given by

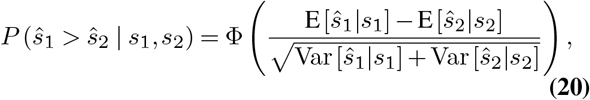

where Φ() is the CDF of the normal distribution. When fitting the choice data to the model, we accounted for potential side (left/right) biases *β*_0_ and lapse rates *λ* in the decision task using

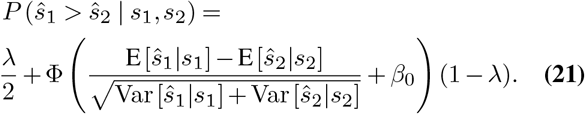

#### Fitting the power-law efficient model

To fit the power-law efficient coding model to the choice data in the “Decision task” we use a hierarchical model. In total we perform 6 separate fits, one for the first and second half of sessions in every context of experiment 1, and one for the first and second half of sessions in experiment 2. Posterior inference of the parameters in the hierarchical models was performed via the Gibbs sampler using the Markov chain Monte Carlo technique implemented in JAGS, assuming flat priors for both the mean and the noise of the estimates. For each model a total of 20,000 samples were drawn from an initial burn-in step and subsequently a total of 10,000 new samples were drawn with three chains (each chain was derived based on a different random number generator engine, and each with a different seed). We applied a thinning of 10 to this final sample, thus resulting in a final set of 1,000 samples for each parameter. We conducted Gelman–Rubin tests for each parameter to confirm convergence of the chains. All latent variables in our Bayesian models had 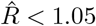, which suggests that all three chains converged to a target posterior distribution. We checked via visual inspection that the posterior population-level distributions of the final Markov chain Monte Carlo chains converged to our assumed parametrizations. We found the *q* and *σ* values for the first and second half of every sessions. The difference between the first and second half of every session is defined as the change Δ*q* and Δ*σ*.

### Behavioral and Statistical analyses

In the “Estimation task”, the observers’ behavioral error on a given trial was computed as the difference between the reported orientation and the presented orientation. The direction of the error was defined as positive if the reported orientation was more oblique than the presented orientation, or negative if vice versa. If the error on any given trial was bigger than 25% of the maximum possible error [90 degrees], we discarded that trial. To make full use of all trials we pooled all sessions in the *K*_*rew*_ context from both experiments for the analysis of the impact of the reward training.

Trials were first sorted into four equally spaced bins of increasing angle from the next cardinal axis. We computed the average bias and variance before the training and after the training in every bin. Next, we computed the average proportional change to the bias and the variance in every bin, as well as over the complete range of trials. To assess the difference between cardinal and oblique trials we compared the most cardinal and the most oblique bin (Fig. 3l, Fig. 4b,c). To assess the effect of context on the variance we first computed the variance for every subject in every bin. Next, we computed the difference between the more oblique and the more cardinal bins (Var(cardinal)-Var(oblique). We employed Bayesian paired t-tests computed with the *BEST* package, implemented in the statistical computing software R, to assess the significance of the differences.

We applied linear mixed effect regression models to investigate the change in absolute value of the bias in the single bins and over the entire range. To this end, we split the data into 6 bins, dependent on the orientation of the stimuli and computed the mean bias as well as the z-scored mean variance for every bin.

The regressions reported for experiments 1 and 2 were computed using the *brm-function* implemented in the statistical computing software R, for linear mixed effect regression models. All of the hierarchical Bayesian (i.e., mixed-effects) regressions we report were specified to have varying subject-specific intercepts and slopes. For each model we used 3-4 chains with at least 1000 samples per chain after burnin. The *P*_*MCMC*_ values reported for these regressions represent the probability of the reported effect being greater or less than zero given the posterior distributions of the fitted model parameters. The *P*_*MCMC*_ values were computed using the posterior population distributions estimated for each parameter and represent the portion of the cumulative density functions that lies above/below 0 (depending on the direction of the predicted effect). We list the mean coefficients, S.D. of the posterior distributions and *P*_*MCMC*_ for the effects of interest in the Results section text, and include the regression tables (with a description of all covariates of interest) for all models the supplementary materials (see Supplementary Tables section).

### Data and code availability

Data and essential code will be made available upon manuscript acceptance.

## ACKNOWLEDGEMENTS

The authors would like to thank Nick Netzer, Michael Woodford, and Alan Stocker for providing helpful feedback on the manuscript text. J.S. gratefully acknowledges support from a Marlene-Porsche Foundation scholarship for his PhD studies. This work was supported by a European Research Council (ERC) starting grant (EN-TRAINER) to R.P. This project has received funding from the European Research Council (ERC) under the European Union’s Horizon 2020 research and innovation program (grant agreement No. 758604). T.A.H received support from the Swiss National Science Foundation (SNSF; grant number 32003B_166566). P.N.T received support from the SNSF (grants 100019_176016 and 10001C_188878)

---

**Figure S1.**
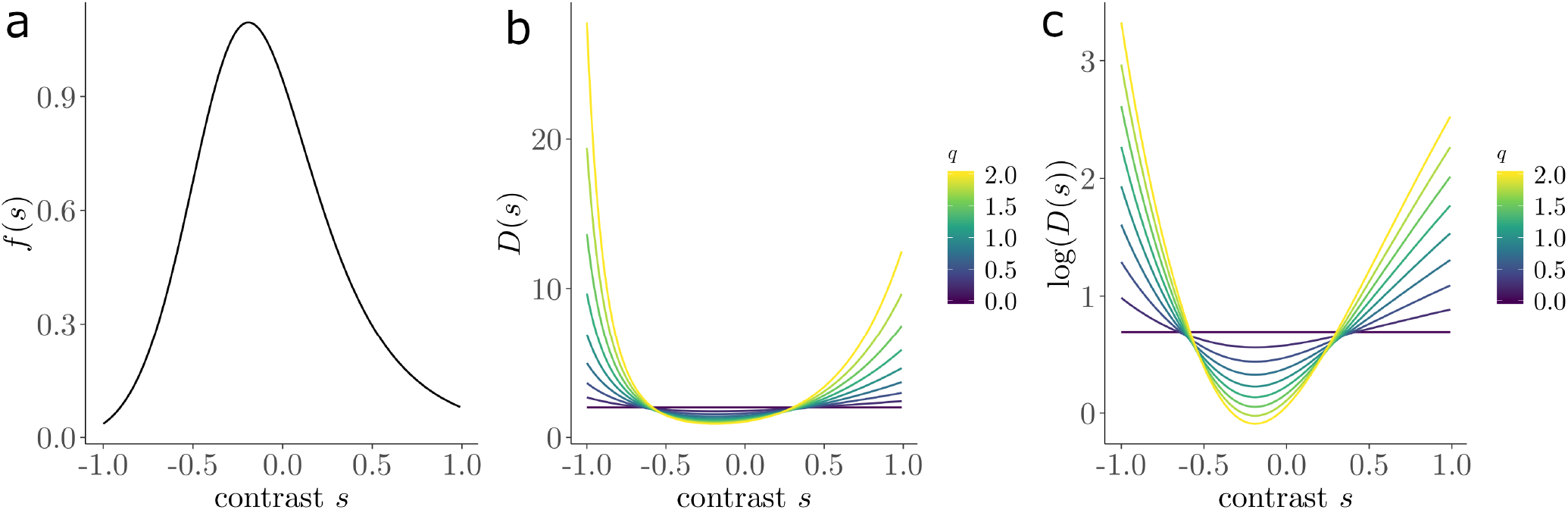
Prior distribution of contrasts in the blowfly and corresponding discriminability thresholds of the power-law efficient codes. **a)** Prior distribution of contrasts *s* that the blowfly experiences in its natural environment. Contrasts that appear more often in the natural environment have higher values on the y-axis. In order to estimate the prior distribution, we followed the procedure from previous work (26) where the stimulus CDF *F* (*s*) was estimated using a 3-parameter Naka-Rushton function

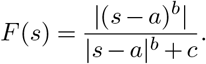

The parameters that best fit the data are *a* = −1.52, *b* = 5.80 and *c* = 7.55 (plotted as the grey dotted line in Fig. 1b). Thus, the prior distribution *f* (*s*) shown in this figure was recovered by integrating over *F* (*s*).

**b, c)** Discriminability thresholds as a function of *s* and power-law value, *q*. Lower values on the y-axis indicate higher discrimination sensitivity. In order to compute the discriminability thresholds *D*(*s*), we used the relationship between *D*(*s*) and Fisher information *J* (*s*) derived in previous work (41). Briefly, assuming that the distribution of stimulus estimations *p*(*ŝ* | *s*) is normally distributed, it is possible to show that the relationship between *D*(*s*) and *J* (*s*) obeys

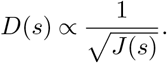

According to the power-law efficient codes, the following relationship between *J* (*s*) and *f* (*s*) holds (see Eq. 9 in Methods)

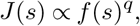

thus,

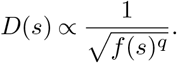

We use this result to generate discriminability threshold predictions as a function of different power *q* values and contrast levels *s* (panel b). Recall that *q* = 2 corresponds to the Infomax solution, and *q* = 4*/*3 to the reward-maximization solution (see Methods). These plots clearly show that higher power *q* values emphasize discriminability for stimuli with higher likelihood of appearing, whereas lower *q* values lead to greater sensitivity for less common stimuli. Panel c shows log(*D*(*s*)) to better visualize the differences in discriminability thresholds.

## Supplementary Tables

**Table S1.** Discrimination accuracy by training context. This table reports the results of a Bayesian hierarchical linear regression on choice performance in the decision task. The regressor *K* is a dummy variable for training type (1 = reward, 0 = accuracy). The significant interaction orientation\**K* indicates that reward training decreased performance in cardinal orientations and increased performance in diagonal orientations, relative to accuracy training.

|  | Estimate | Est.Error | Pmcmc |
| --- | --- | --- | --- |
| Intercept | 85.62 | 1.24 | 0.0001 |
| orientation ($s$) | -0.48 | 0.07 | 0.0001 |
| $K$ | -4.55 | 1.66 | 0.0053 |
| orientation*$K$ | 0.26 | 0.11 | 0.0105 |

**Table S2.** Changes in estimation bias following reward training. This table reports the results of a Bayesian hierarchical linear regression on estimation bias after training in the reward context, *K*_rew_.The regressors *s* and *s*^2^ represent the linear and quadratic effects of orientation angle. The regressors controlling for session effects are not displayed for conciseness. The reward training context leads to a significant reduction in estimation bias as predicted by the fitness maximization hypothesis.

|  | Estimate | Est.Error | Pmcmc |
| --- | --- | --- | --- |
| Intercept | 5.78 | 0.34 | 0.0001 |
| Training | -0.37 | 0.18 | 0.0230 |
| $s$ | -1.23 | 0.11 | 0.0001 |
| $s^2$ | -1.21 | 0.13 | 0.0001 |

**Table S3.**
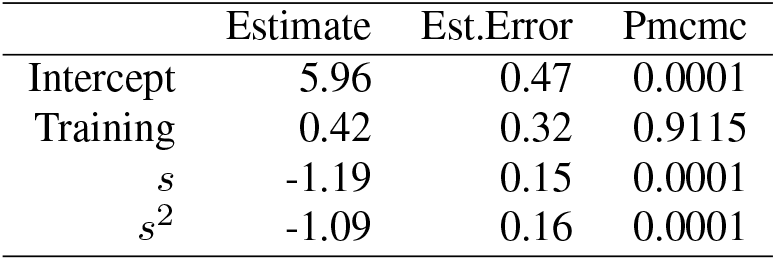
Changes in estimation bias following accuracy training. This table reports the results of a Bayesian hierarchical linear regression on estimation bias after training in the accuracy context, *K*_acc_. The regressors *s* and *s*^2^ represent the linear and quadratic effects of orientation angle. The regressors controlling for session effects are not displayed for conciseness. There is no significant effect of accuracy training on estimation bias. Neither the fitness maximization nor information maximization schemes predict a change in bias following this type of training. The *P*_*MCMC*_ values were computed using the posterior population distributions estimated for each parameter and represent the portion of the cumulative density functions that lies above/below 0 depending on the sign of the effect. Estimation error is quantified as the standard deviation of the posterior distribution of estimates.

**Table S4.** Changes in estimation bias following reward versus accuracy training. This table reports the results of a Bayesian hierarchical linear regression on estimation bias after training in the reward versus accuracy contexts. The regressors *s* and *s*^2^ represent the linear and quadratic effects of orientation angle. The regressor *K* is a dummy variable for training type (1 = reward). The regressors controlling for session effects are not displayed for conciseness. The significant Training\**K* interaction effect indicates that reward training changes estimation bias more than accuracy training. This interaction effect is predicted by the fitness maximization hypothesis, but is inconsistent with information maximization coding schemes. The *P*_*MCMC*_ values were computed using the posterior population distributions estimated for each parameter and represent the portion of the cumulative density functions that lies above/below 0 depending on the sign of the effect. Estimation error is quantified as the standard deviation of the posterior distribution of estimates.

|  | Estimate | Est.Error | Pmcmc |
| --- | --- | --- | --- |
| Intercept | 6.16 | 0.39 | 0.0000 |
| Training | 0.40 | 0.27 | 0.0712 |
| $K$ | -0.36 | 0.34 | 0.1482 |
| $s$ | -1.22 | 0.10 | 0.0000 |
| $s^2$ | -1.21 | 0.12 | 0.0000 |
| Training*$K$ | -0.79 | 0.32 | 0.0085 |

**Table S5.** Changes in the variance in estimation accuracy following reward training. This table reports the results of a Bayesian hierarchical linear regression on the variance in estimation accuracy after training in the reward contexts. The regressor *s* represents the linear effect of orientation angle. The significant interaction Training*s indicates a decrease in variance for cardinal orientations and an increase in variance for diagonal orientations. The *P*_*MCMC*_ values were computed using the posterior population distributions estimated for each parameter and represent the portion of the cumulative density functions that lies above/below 0 depending on the sign of the effect. Estimation error is quantified as the standard deviation of the posterior distribution of estimates.

|  | Estimate | Est.Error | Pmcmc |
| --- | --- | --- | --- |
| Intercept | 65.25 | 2.58 | 0.0001 |
| Training | -2.19 | 1.49 | 0.0689 |
| $s$ | 11.63 | 1.76 | 0.0001 |
| Training*$s$ | -2.70 | 1.49 | 0.0347 |

**Table S6.**
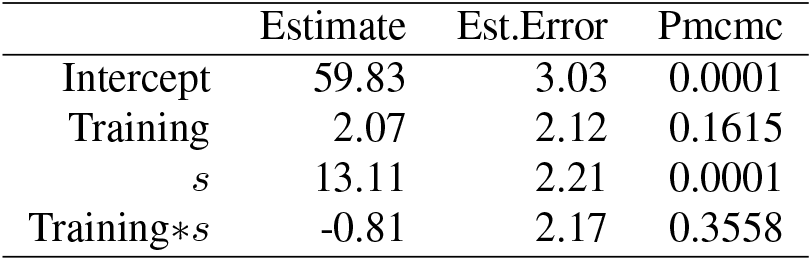
Changes in the variance in estimation accuracy following accuracy training. This table reports the results of a Bayesian hierarchical linear regression on the variance in estimation accuracy after training in the accuracy context. The regressor *s* represents the linear effect of orientation angle. The accuracy training context does not lead to significant reduction in variance. The *P*_*MCMC*_ values were computed using the posterior population distributions estimated for each parameter and represent the portion of the cumulative density functions that lies above/below 0 depending on the sign of the effect. Estimation error is quantified as the standard deviation of the posterior distribution of estimates.

**Table S7.** Effects of changes in the power-law parameter and encoding noise on changes in estimation bias. This table reports the results of a Bayesian hierarchical linear regression on the effects of the change in the power-law parameter (Δ*q*) and encoding noise (Δ*σ*) on changes in estimation bias. The significant interaction Δ*q * K*_*rew*_ indicates that changes in *q* induced by reward training were associated with changes in estimation bias. On the other hand, changes in *q* were not related to changes in estimation bias following *K*_*acc*_ training (Δ*q * K*_*acc*_). The *P*_*MCMC*_ values were computed using the posterior population distributions estimated for each parameter and represent the portion of the cumulative density functions that lies above/below 0 depending on the sign of the effect. Estimation error is quantified as the standard deviation of the posterior distribution of estimates.

|  | Estimate | Est.Error | Pmcmc |
| --- | --- | --- | --- |
| Intercept | 1.46 | 8.54 | 0.4297 |
| $\Delta q$ | 14.42 | 10.52 | 0.0835 |
| $\Delta\sigma$ | -10.85 | 13.15 | 0.2040 |
| $K_{acc}$ | -14.50 | 20.04 | 0.2255 |
| $K_{rew}$ | -14.69 | 9.04 | 0.0525 |
| $\Delta q * \Delta\sigma$ | 5.25 | 12.16 | 0.3220 |
| $\Delta q * K_{acc}$ | -10.44 | 16.43 | 0.2592 |
| $\Delta q * K_{rew}$ | -24.67 | 11.01 | 0.0152 |
| $\Delta\sigma * K_{acc}$ | -15.05 | 25.39 | 0.2730 |
| $\Delta\sigma * K_{rew}$ | 18.79 | 13.44 | 0.0830 |
| $\Delta q * \Delta\sigma * K_{acc}$ | 8.27 | 19.49 | 0.3405 |
| $\Delta q * \Delta\sigma * K_{rew}$ | 0.22 | 12.38 | 0.4970 |

**Table S8.**
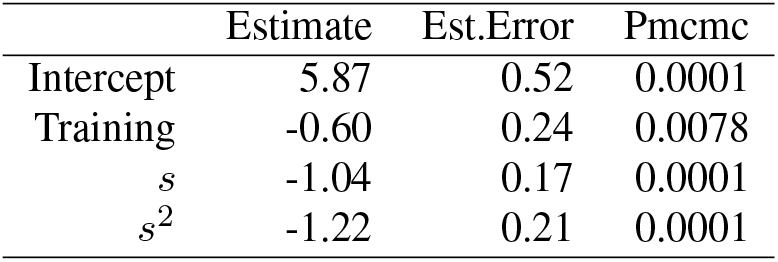
Changes in the estimation bias for the trained location in Experiment 2. This table reports the results of a Bayesian hierarchical linear regression on the estimation bias for the location trained in the reward context. The regressors *s* and *s*^2^ represent the linear and quadratic effects of orientation angle. The regressors controlling for session effects are not displayed for conciseness. The bias in the trained condition is significantly lower after training.The *P*_*MCMC*_ values were computed using the posterior population distributions estimated for each parameter and represent the portion of the cumulative density functions that lies above/below 0 depending on the sign of the effect. Estimation error is quantified as the standard deviation of the posterior distribution of estimates.

|  | Estimate | Est.Error | Pmcmc |
| --- | --- | --- | --- |
| Intercept | 5.87 | 0.52 | 0.0001 |
| Training | -0.60 | 0.24 | 0.0078 |
| $s$ | -1.04 | 0.17 | 0.0001 |
| $s^2$ | -1.22 | 0.21 | 0.0001 |

**Table S9.**
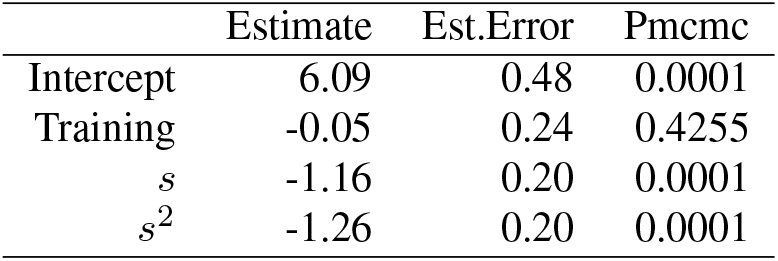
Changes in the estimation bias for the untrained location in experiment 2. This table reports the results of a Bayesian hierarchical linear regression on the estimation bias for the location untrained in the reward condition. The regressors *s* and *s*^2^ represent the linear and quadratic effects of orientation angle. The regressors controlling for session effects are not displayed for conciseness. The bias in the untrained condition is not significantly different after training. The *P*_*MCMC*_ values were computed using the posterior population distributions estimated for each parameter and represent the portion of the cumulative density functions that lies above/below 0 depending on the sign of the effect. Estimation error is quantified as the standard deviation of the posterior distribution of estimates.

**Table S10.** Changes in the estimation bias for the trained versus untrained locations in experiment 2. This table reports the results of a Bayesian hierarchical linear regression on the estimation bias for the trained versus untrained locations in the reward condition. The regressors *s* and *s*^2^ represent the linear and quadratic effects of orientation angle. Location is a dummy variable indicating whether the stimuli appeared in the trained or untrained locations (1 = trained, 0 = untrained). The dummy variables controlling for session effects that are not displayed for conciseness. The bias change in the trained condition is significantly greater than the untrained location (Training*Location interaction). The *P*_*MCMC*_ values were computed using the posterior population distributions estimated for each parameter and represent the portion of the cumulative density functions that lies above/below 0 depending on the sign of the effect. Estimation error is quantified as the standard deviation of the posterior distribution of estimates.

|  | Estimate | Est.Error | Pmcmc |
| --- | --- | --- | --- |
| Intercept | 6.10 | 0.49 | 0.0001 |
| Training | -0.06 | 0.24 | 0.4065 |
| Location | -0.15 | 0.24 | 0.2645 |
| $s$ | -1.11 | 0.16 | 0.0001 |
| $s^2$ | -1.25 | 0.19 | 0.0001 |
| Training*Location | -0.55 | 0.32 | 0.0425 |

